# The multiscale distribution of radiation-induced DNA damage and its impact on local genome structure

**DOI:** 10.1101/2025.08.25.672161

**Authors:** Heng Li, Tianchun Xue, Rachel Patton McCord

## Abstract

The three-dimensional genome structure is critical for the regulation of gene expression and repair of DNA damage. While previous work has characterized genome-wide sites of DNA damage caused by etoposide or nucleases, the distribution of double-strand breaks (DSBs) caused by external radiation and how these interact with the 3D genome organization is less well understood. Here, we measure the genomic landscape of radiation-induced DNA damage using END-seq in fibroblasts and lymphoblasts after exposure to 5 Gy X-rays. We identify frequently broken regions and investigate the 3D genome properties around these breaks with Hi-C data. We observe that the distribution of robust breaks correlates with transcriptional and chromatin features of the genome. Transcriptionally active and decondensed regions, such as chromosome 19, the A compartment, and topologically associating domain (TAD) boundaries, show pronounced break probability. We also find evidence of DSB-induced loop formation in the vicinity of frequent radiation-induced breaks. Our data reveal that pre-existing 3D genome architecture influences the distribution of radiation-induced DSBs and that these breaks reshape local chromatin landscapes.

## Introduction

Ionizing radiation (IR) is a potent genotoxic agent that can induce various types of DNA damage, most notably double-strand breaks (DSBs), compromising genomic integrity and contributing to mutagenesis, carcinogenesis and cell death. Thus, appropriate and faithful repair of DSBs is essential for maintaining genomic integrity. Previous research has reported numerous ways that DNA damage and repair can be influenced by the packaging of DNA into the nucleus in the 3D genome structure^1^, indicating that the local genomic landscape might influence the recognition of DNA damage and the selection of DNA repair pathways^2,3^.

In eukaryotes, DNA is folded into the nucleus in a multi-scale packaging called three-dimensional (3D) structure. With the advent of genome-wide chromosome conformation capture technology Hi-C, chromatin is generally divided into two spatial compartments by principal components analysis (PCA) of the Hi-C contact matrix. A and B compartments correspond to the euchromatic, open, and transcriptionally active regions and heterochromatic, condensed, and transcriptionally inactive regions, respectively^4–6^. Additionally, topologically associating domains (TADs) and loops are organized by CCCTC-binding factor (CTCF) and cohesin-driven loop extrusion to foster appropriate gene regulation^7^. Emerging evidence suggests that the spatial organization of the genome within nucleus plays a crucial role in the detection, signaling and repair of DSBs. TAD structure has been reported to confine the spatial propagation of phosphorylated H2AX (γH2AX), a marker of DNA damage^8^. Studies on restriction enzyme-induced DSBs have shown that DSBs at transcriptionally active loci are physically relocated toward the nuclear periphery, showing shifts in compartmentalization, and that cohesin-mediated loop extrusion occurs at DSBs and contributes to their clustering^9–11^. Nuclear actin polymerization can facilitate these compartment switches and contribute to the DSB clustering and repair^12^. Additional studies have reported a strengthening of TAD boundaries after site specific nuclease damage as well as externally induced damage by UV and X-ray irradiation^10,13,14^. Given the influence of 3D genome structure after DNA damage, it suggests a vital function of 3D genome organization in DNA damage and repair.

IR can induce DNA damage both through direct interaction with DNA and indirectly via the ionization of water molecules, leading to the generation of free radicals and reactive oxygen species (ROS) that damage DNA, lipids and proteins. These processes produce a wide variety of DNA lesions at stochastic positions across the genome^15^. However, early studies showed that chromosomal aberrations may tend to arise in euchromatin^16–18^. Euchromatin is generally accessible with lower nucleosome density than heterochromatin. Cells with lower nucleosome occupancy or partially depleted histone proteins are sensitive to IR-induced DNA damage^19,20^. In line with the given results, imaging-based foci assays and computational simulation models have reported that condensed chromatin or heterochromatin protects against DNA damage by IR^21–24^. These results indicate that DSB susceptibility may be influenced by local chromatin accessibility. Accordingly, it is important to understand whether the 3D genome organization impacts the induction of DNA damage by IR and whether IR-induced DNA breaks alter the local genome architecture.

Numerous sequencing-based techniques such as BLISS, BLESS, DSBCapture, Damage-seq and END-seq have been previously used to capture the genome-wide distribution of DSBs caused by agents that tend to act at particular genomic locations^25–29^. For example, END-seq experiments revealed that DSBs pile up near TAD boundaries after the inhibition of topoisomerase with etoposide^30^. Despite the stochastic nature of radiation-induced DNA breaks, Kaya *et al.* measured the distribution of UV-induced damage at TAD boundaries using Damage-seq^13^, and Brambilla *et al.* reported that the distribution of IR-induced DSBs correlated with nucleosome occupancy, as determined by BLISS in mouse cells^20^. Here, we quantify X-ray induced DSBs using END-seq, where, in contrast to the techniques employed in the studies mentioned above, cells are embedded in an agarose gel plug without fixation. We examine two different human cell types and use a multiscale analysis to identify sites of frequently occurring DSBs and to compare their distribution to other features and 3D genome structure. We found that while the number of robust breaks is correlated with the size and nuclear position of each chromosome, the break probability is determined by the DNA accessibility at each genome structure length scale (chromosome, compartment, TAD) in both cell types. By examining the alterations in 3D genome organization at 30 min and 24 h after IR, we found stripe patterns at sites of frequent IR-induced breaks in fibroblasts, suggesting these breaks nucleate increased loop extrusion as has been observed in the AsiSI-model^10^.

## Results

### Radiation-induced robust breaks are identified by END-seq

To characterize the genomic distribution of radiation-induced DSBs, we used human fibroblasts (BJ-5ta) and lymphoblasts (GM12878). These cell types have notably different nucleus geometries, chromosome territory positions, and inherent radiosensitivity^31–34^, allowing us to determine whether we can detect meaningful patterns of DNA damage that correlate with these nucleus architecture and genome structure differences. We have also previously characterized the 3D genome response to irradiation in these cell types at 30 min and 24 h after exposure to 5 Gy X-rays^35^. About 100 - 150 DSBs are generated in each cell exposed to 5 Gy X-ray^36^. To accurately map the position of radiation-induced DSBs, we used the sensitive and quantitative method END-seq^37^. BJ-5ta and GM12878 cells were collected immediately after X-ray exposure and processed following the END-seq protocol (**Figure 1a**).

**Figure 1:**
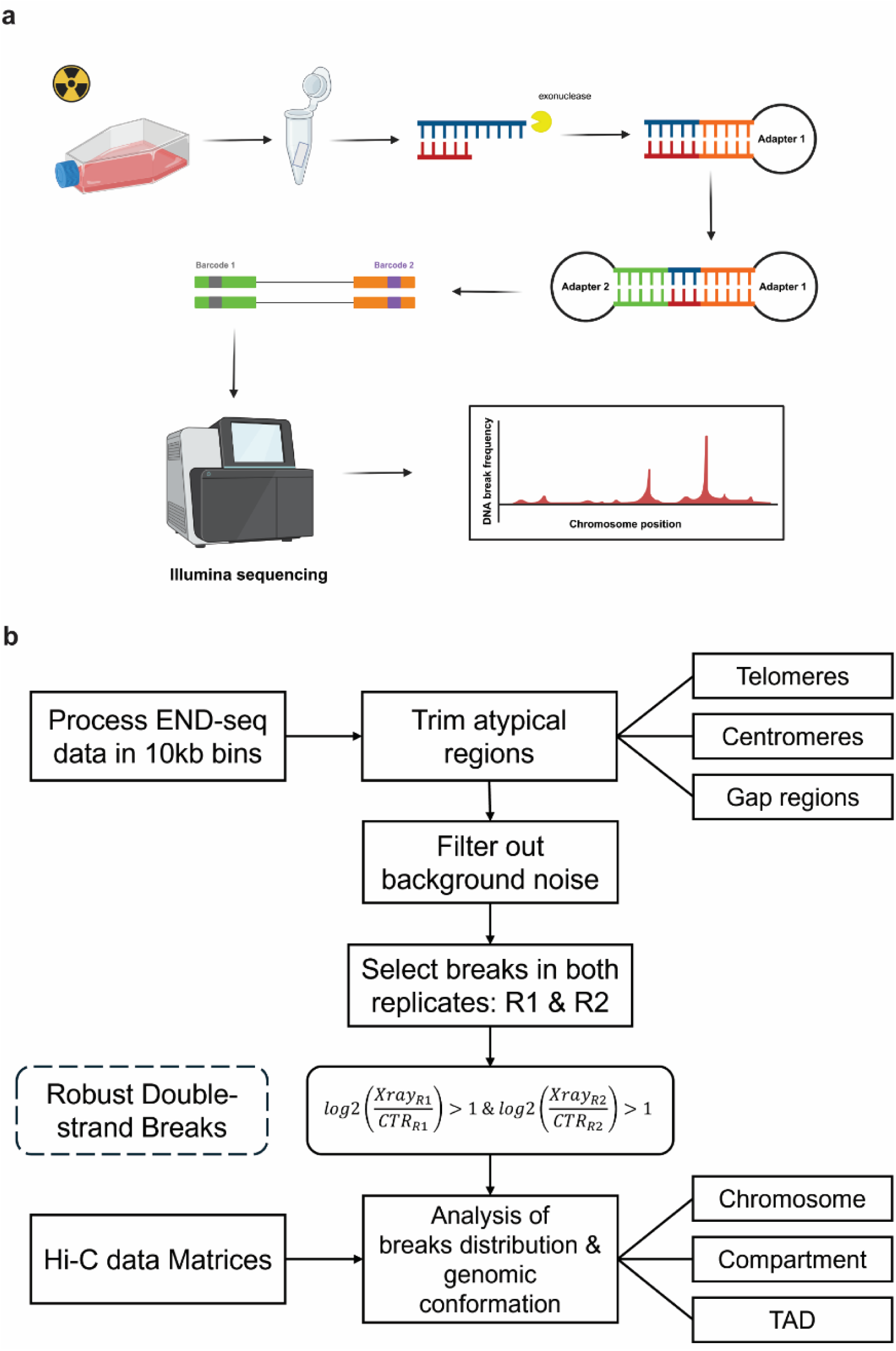
Overview and analytics of END-seq. **a.** Schematic of the END-seq workflow. Cells are irradiated with 5 Gy X-rays and collected and embedded into an agarose gel plug. Damaged ends are processed and ligated with adapters. Amplified PCR library is ready to sequence with the Illumina NovaSeq platform. **b.** Schematic illustrates the process of analyzing break frequency overlapping with differential 3D genomic landscape.

As a positive control for the END-seq method, we used the AsiSI (DIvA) system^38^, which is induced by 4OHT (4-hydroxytamoxifen) and digests the DNA at an 8 bp target sequence (GCGATCGC) across the genome, generating DSBs (**Supplementary Figure 1a**). We detected sharp and intense peaks precisely aligned at predicted AsiSI sites after a 4 h 4OHT treatment in U2OS-ER-AsiSI and BJ-5ta-ER-AsiSI cells, while negligible signal was observed in undamaged conditions (**Supplementary Figure 1b**). Additionally, we identified cell-type specific END-seq peaks at certain predicted AsiSI sites, indicating that the action of the restriction enzyme AsiSI is dependent on the accessibility of genomic landscape in the given cell line (**Supplementary Figure 1c**). Relatedly, as previously reported, we observe that a subset of predicted AsiSI sites show no END-seq peaks^2^ (**Supplementary Figure 2a**). Overall, END-seq detects a similar set of AsiSI induced breaks when compared to the previously published BLESS method^39^ (**Supplementary Figure 2b-c**). These results suggest that END-seq has the capacity to capture double-strand breaks, and also to show differential peaks in different cell types.

To investigate the distribution of radiation-induced DSBs genome wide, we devised an analysis pipeline using replicates of END-seq signal across the genome before and after X-ray irradiation to determine the most frequently damaged regions (**Figure 1b**). First, we observed that extremely strong signals were detected at telomere and centromere regions, where repetitive sequences are enriched and prone to fragility^40–42^. To eliminate the effect of non-IR-induced DNA breaks, we excluded telomeres, centromeres and gap regions from our further analyses. Then, genome-wide END-seq signals were binned (50 kb) and normalized across conditions, and regions showing consistent enrichment in both replicates were selected based on a threshold of log_2_(X-ray/Control) > 1 (**Supplementary Figure 3a**). Regions passing this criterion were identified as ‘robust breaks’ in each cell line and were used for all subsequent analyses of radiation-induced DNA damage distribution (**Supplementary Figure 3b**). Each robust break bin is associated with a “break frequency”, which is the sum of END-seq reads in that bin. Unlike AsiSI-induced DSBs, where break frequency is very high for the limited number of predicted sites that are both accessible and cleavable but then drastically decreases, radiation-induced DSBs show a relatively homogeneous break frequency (**Supplementary Figure 3c, d**). This indicates that there are many break sites that experience a similar extent of DNA damage across the cell population.

### Radiation-induced DSBs are determined by chromosome features

Firstly, we detected the number of robust breaks in each chromosome and observed that, in both fibroblasts and lymphoblasts, chromosome size was generally correlated with the total number of breaks (Spearman’s rank correlation (SCC) of 0.96 and 0.94 in fibroblasts and lymphoblasts, respectively) (**Figure 2a**). We next considered whether the divergent radial positioning of chromosomes in fibroblasts compared to lymphoblasts^33,34^ would have an impact on whole chromosome break counts. We previously predicted, using 3D reconstructions of Hi-C data and DNA damage simulations, that differences in chromosome arrangements between these cell types would result in different amounts and rates of initial repair of DSBs^32^. We performed correlation analysis on the relative spatial distance to the nuclear center^31^ for each individual chromosome in fibroblasts and lymphoblasts. The number of robust breaks is positively correlated with the spatial arrangement of each chromosome within the nucleus, with chromosomes located closer to the center of the nucleus exhibiting significantly less DNA damage (**Figure 2b**).

**Figure 2:**
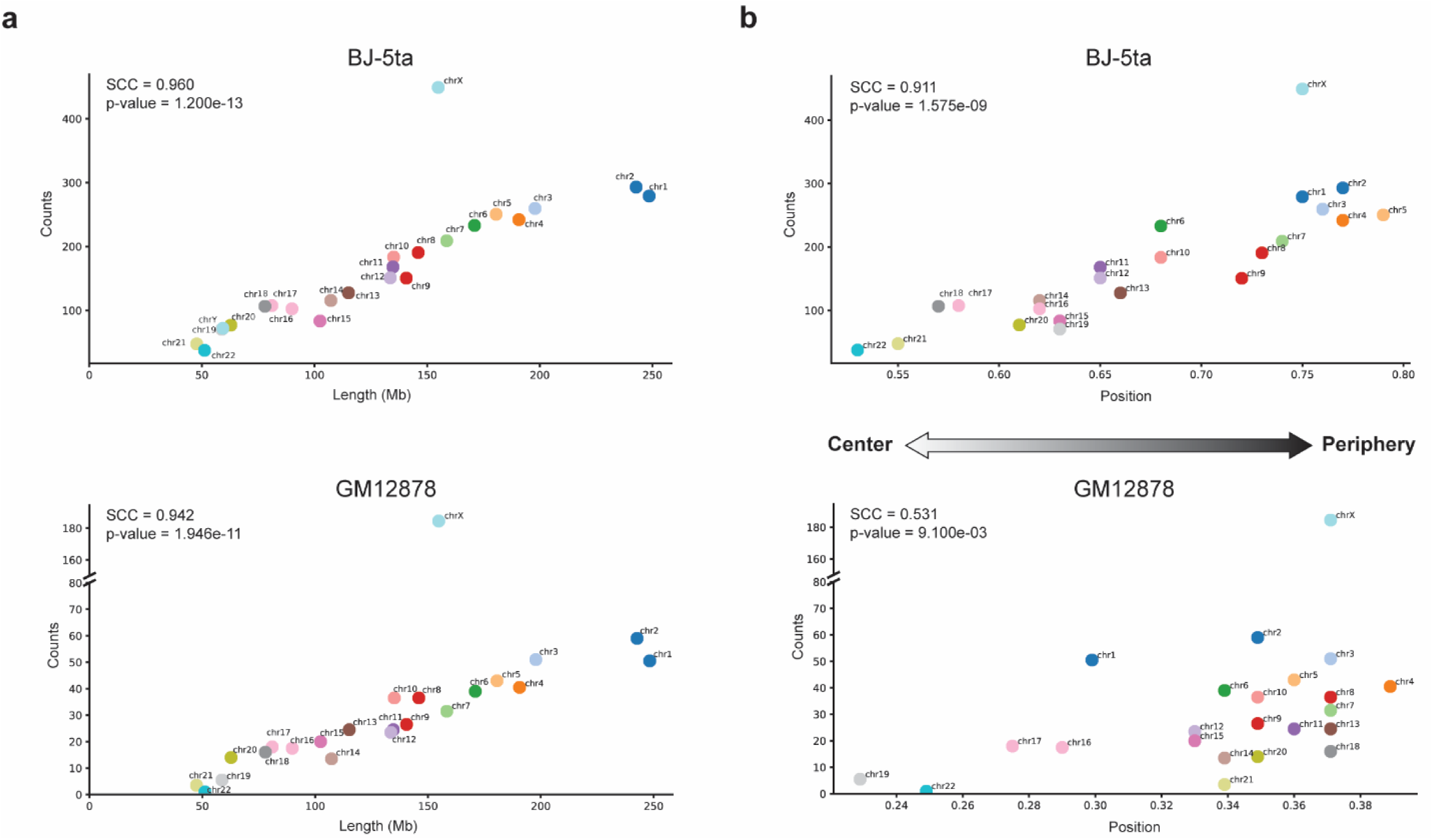
Chromosomal-level patterns of robust DNA break counts. **a.** Scatter plots showing the relationship between the length of chromosomes and the number of robust breaks in BJ-5ta (top) and GM12878 (bottom) cells. **b.** Scatter plots exhibiting the relationship between the position of chromosomes within the nucleus and the number of robust breaks in BJ-5ta (top) and GM12878 (bottom) cells. All p-values are from Spearman correlation test as indicated in figures.

We found that chromosome X is an outlier to the trend described above. Though its size is similar to chromosome 7 and chromosome 8, it exhibited an extraordinary number of breaks, so we examined the location of these breaks across the whole chromosome in detail. First, we note that the BJ-5ta cells are male and thus have only a single X chromosome with no inactivation. For these cells, the robust break sites were distributed evenly across the chromosome. In contrast to the fibroblasts, the female GM12878 lymphoblast cell line showed an asymmetrical distribution of DNA breaks, with a significantly higher incidence on the q arm compared to the p arm (**Figure 3a**). These female cells have one active X and one inactivated X chromosome. Interestingly, despite a higher total number of robust breaks on the q arm, the total break frequency was higher on the p arm, indicating that p-arm breaks are more recurrent and concentrated in specific regions (**Figure 3b**). We hypothesized that this difference between chromosome arms in the female cell line could reflect the known uneven pattern of X chromosome inactivation, in which there are more genes that escape inactivation on the p arm of the X chromosome^43,44^. Indeed, when we analyzed published RNA-seq data for GM12878 cells, we found that there is a higher gene expression density on the p arm, as reflected by the sum of RNA-seq counts normalized by length^45,46^ (**Figure 3c**). This suggests that the p arm of the inactive X chromosome more frequently experiences breaks in a few locations across the population, associated with the few highly expressed genes on this arm. Meanwhile, the q arm experiences more independent break locations, but each at lower frequency.

**Figure 3:**
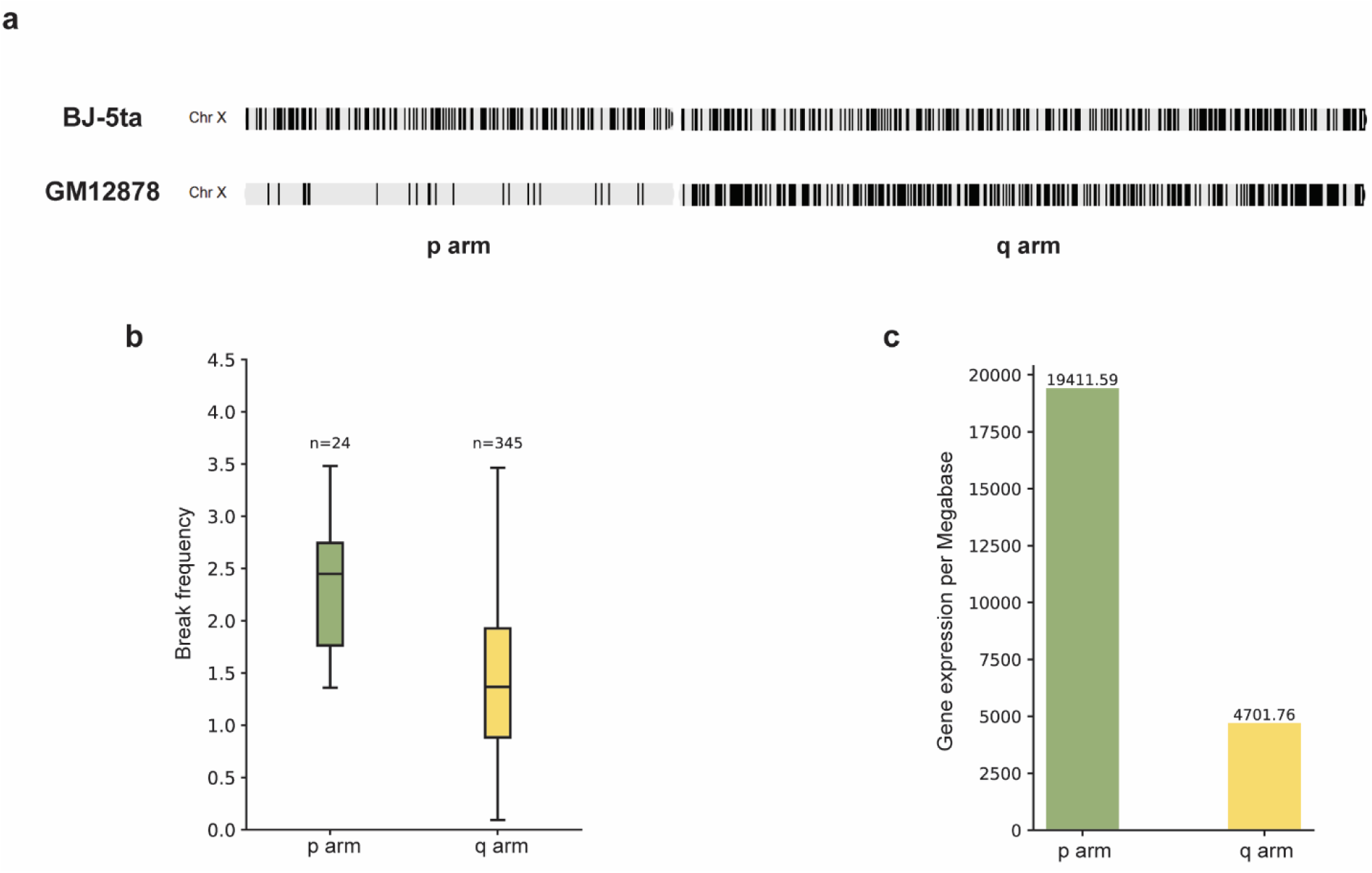
Asymmetric breaks and expression along chromosome X in GM12878. **a.** The distribution of robust breaks in chromosome X after IR BJ-5ta (top) and GM12878 (bottom) cells. Each tick mark represents a 50 kb bin classified as a robust break superimposed on the chrX diagram. **b.** Boxplots representing the break frequency in 50 kb windows on the p and q arm of chromosome X in GM12878 cells. **c.** Histogram showing the sum of RNA-seq signal normalized by chromosome arm length for the p and q arm of chromosome X in GM12878 cells.

To further assess the impact of gene expression and gene density on the susceptibility to DSBs, we examined break frequency of genomic regions on autosomes. Of note, a significant difference of break frequency was observed between two autosomes of similar size, chromosome 18 and chromosome 19. Though the total number of robust break locations was higher in chromosome 18, chromosome 19 showed a higher break frequency compared to chromosome 18 in both fibroblasts and lymphoblasts, corresponding to its higher gene density (**Figure 4, Supplementary Figure 4**. These results indicate another association between break frequency and higher gene density and elevated gene expression activity, suggesting a potential link between transcriptional activity and radiation-induced genomic instability.

**Figure 4:**
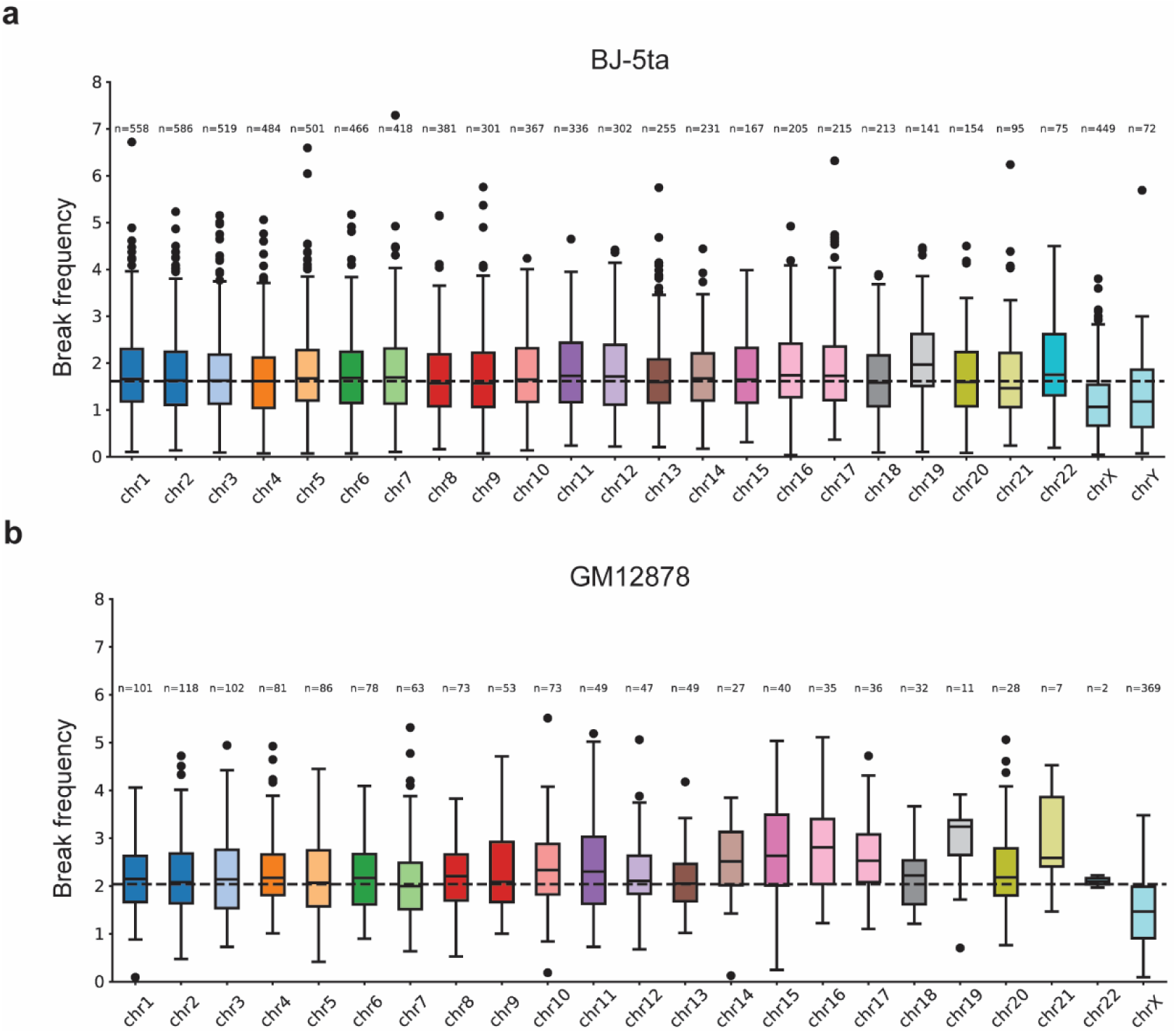
Chromosomal-level patterns of robust DNA breaks by break frequency. **a, b.** Boxplots representing the break frequency in each 50 kb window for all chromosomes in replicate 1 in BJ-5ta (**a**) and GM12878 (**b**) cells. Boxes are colored by chromosomes. Dashed lines represent the median value across all chromosomes. Boxes show 25^th^ percentile, median, and 75^th^ percentile with whiskers showing up to 1.5 times the interquartile range. Numbers of bins on each chromosome are shown above each box.

### Open genomic landscapes experience more frequent radiation-induced DNA damage

The intensity of radiation-induced DNA damage across different genomic regions is influenced by chromatin compaction, as shown by both quantification of DSB signal (H2AX immunofluorescence) and computational modelling studies^21,24^. To characterize the distribution of robust DNA damage sites across different layers of genomic organization, we compared break locations with our previously generated Hi-C datasets in fibroblasts and lymphoblasts exposed to 5 Gy X-rays at 30 min and 24 h post-irradiation^14^.

Hi-C studies have shown that the genome is broadly spatially organized into two compartments, A and B, which tend to have different accessibility, transcription levels, and histone modifications^47^. We next examined whether genomic regions in these distinct chromatin landscapes display differential levels of DNA damage following radiation exposure. To determine the distribution of robust breaks into each compartment, 250 kb binned matrices were identified as A or B compartments by the first eigenvector from PCA. We observed that the break frequency in A compartment was significantly higher than in B compartment in both fibroblasts and lymphoblasts. As with the contrast between chromosome 18 and 19, however, enrichment analysis showed that the number of robust break locations was higher within B compartments. (**Figure 5a, Supplementary Figure 5a**). These results were also consistent with our observations described above of higher break frequency at fewer robust break locations for certain regions of chromosome X.

**Figure 5:**
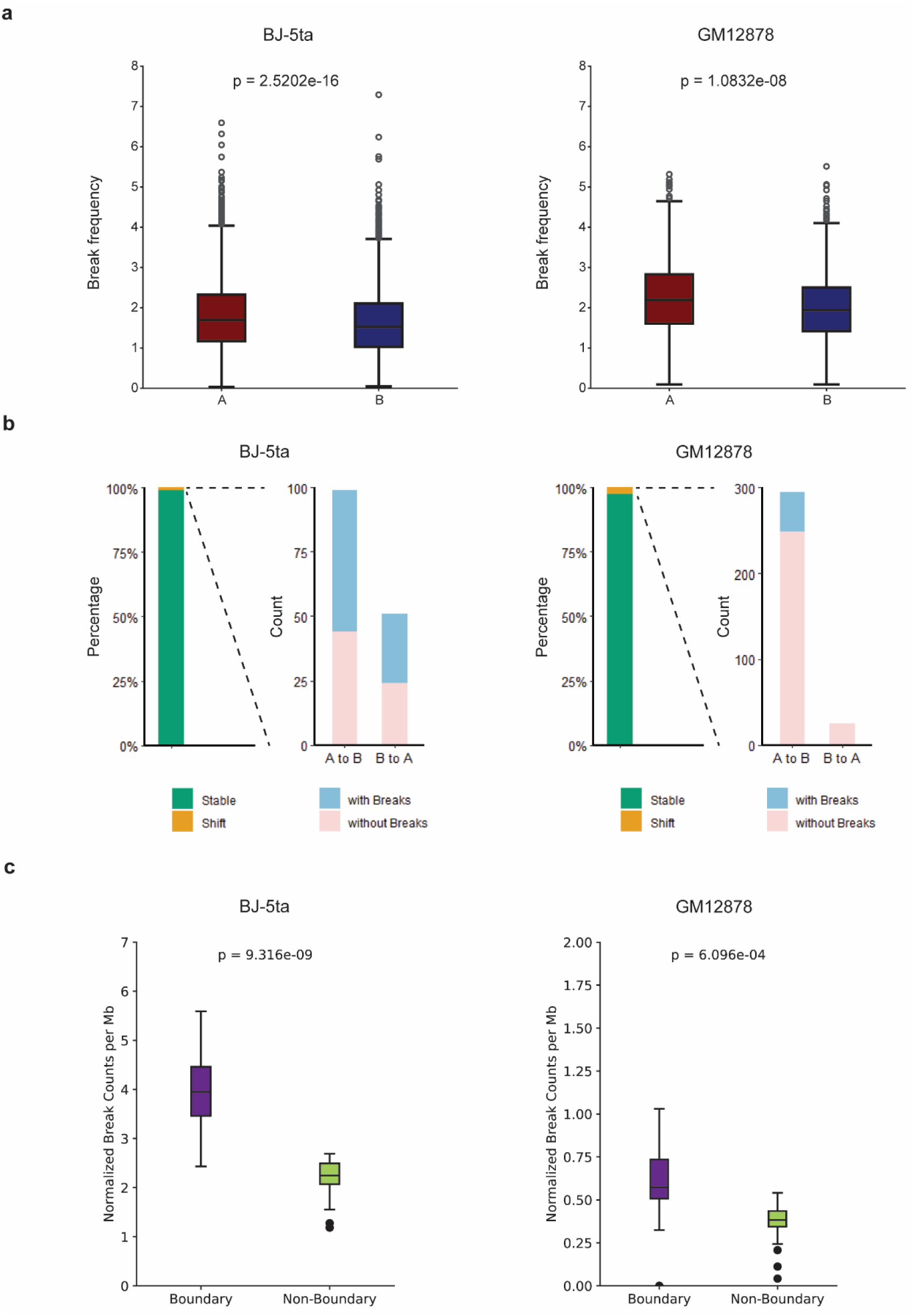
Distribution of DNA breaks relative to 3D genome organization layers. **a.** Boxplots representing the break frequency in each 50 kb window for A (red) and B (blue) compartments in BJ-5ta (left) and GM12878 (right) cells in replicate 1. **b.** Bar plots showing the counts of 250 kb bins with (sky blue) or without (rose) robust breaks within compartments that shifted between control and 30 min post-irradiation, including A to B and B to A, in BJ-5ta (left) and GM12878 (right) cells. **c.** Boxplots representing the counts of robust breaks per megabase within TAD boundary regions (dark purple) or non-boundary regions (lime) in BJ-5ta (left) and GM12878 (right) cells. Boxes show 25^th^ percentile, median, and 75^th^ percentile with whiskers showing up to 1.5 times the interquartile range. Numbers of bins on each chromosome are shown above each box. P-value calculated using Mann-Whitney U test.

Compartment switches in response to changes in conditions have been identified as one of the chromosome reorganizations which may influence biological function, including gene regulation, DNA damage and repair^6^. Previous studies reported that compartments with AsiSI-induced DSBs were partially switched by either nuclear actin polymerization or in PER2-dependent manner^12,48^. However, only a few compartment shifts were observed after exposure to IR^14^. To investigate whether compartment flips occur at sites of frequent DNA breaks after IR, we identified switched compartments at 30 min and 24 h after IR in both BJ-5ta and GM12878 and calculated the proportion of the compartments containing robust breaks. Although less than 1% compartments flipped in BJ-5ta at 30 min post-IR, we observed a marginal majority harboring robust breaks, suggesting that these sites of frequent DNA breaks might give rise to the reconfiguration of compartmentalization as observed in enzyme induced DNA damage^12^ (**Figure 5b, Supplementary Figure 5b**). In contrast to this effect in fibroblasts, GM12878 cells showed only a few shifted compartments that included strong DNA breaks. This might be due to a distinct response to DNA damage in different cell types and stages of the cell cycle.

Topologically associating domains (TADs) are the fundamental unit of 3D genome organization, in which regulatory elements and their target genes are organized, and can be identified as interacting squares along the diagonal in Hi-C data. According to the loop extrusion model, TAD formation is mediated by the cohesin complex, which extrudes DNA into a loop until it is halted by boundary elements, such as CTCF^7,49,50^. Similar to the TAD boundary strengthening which we previously observed after IR, an increase in boundary strength was revealed following UV exposure, and DNA repair activity was enriched at most insulated TAD boundaries^13,14^. TAD boundaries are enriched for insulator proteins, housekeeping genes and transfer RNA genes, and consequently boundary regions exhibit higher chromatin accessibility compared to the rest of DNA^7,51,52^. Indeed, quantitative analysis revealed a significant enrichment of DSBs at TAD boundaries in either endogenous or etoposide-induced DSBs^53^. Therefore, we examined the abundance of DNA damage at boundary sites after IR. To measure the probability of DNA breaks at TAD boundaries, we examined the number of robust breaks within TAD boundaries and non-boundary regions. Notably, break counts were significantly higher at TAD boundaries than non-boundaries, suggesting that boundary regions show pronounced vulnerability to radiation (**Figure 5c**).

Overall, our results suggest that DSB probability is impacted by DNA accessibility in a multiscale genome organization after IR.

### Loop extrusion signatures are identified at robust DSBs

Fundamental genomic structures, including TADs and loops, are established and maintained through cohesin-dependent dynamics. While cohesin was initially identified to remain sister chromatids together from the beginning of S phase, there is strong evidence showing the role of cohesin in maintaining genome stability and preventing the mobility of DSBs from mis-joining of distant DSB ends^54^. Notably, the enrichment of cohesin at DSBs locally leads to the increase of DNA looping in both G1 and G2 phase after AsiSI-induced or CRISPR-Cas9 induced DSBs, which may facilitate the propagation of γH2AX signal and the homology search^10,55,56^. To investigate whether loop extrusion can be observed at IR-induced DSBs, we conducted a comparative analysis of Hi-C contact maps generated before and after IR exposure (**Figure 6**). Our analysis focused on cis contact frequencies in the vicinity of robust DSBs to assess local chromatin architecture changes potentially affected by DNA damage. Differential Hi-C contact maps (24 h - Control) revealed evident stripe-like patterns flanking DSB sites in both fibroblasts and lymphoblasts, indicating the loop extrusion activity. In contrast, only subtle changes were observed in the differential maps (30 min - Control), suggesting that local chromatin reorganization around robust DSBs is limited at early time points.

**Figure 6.**
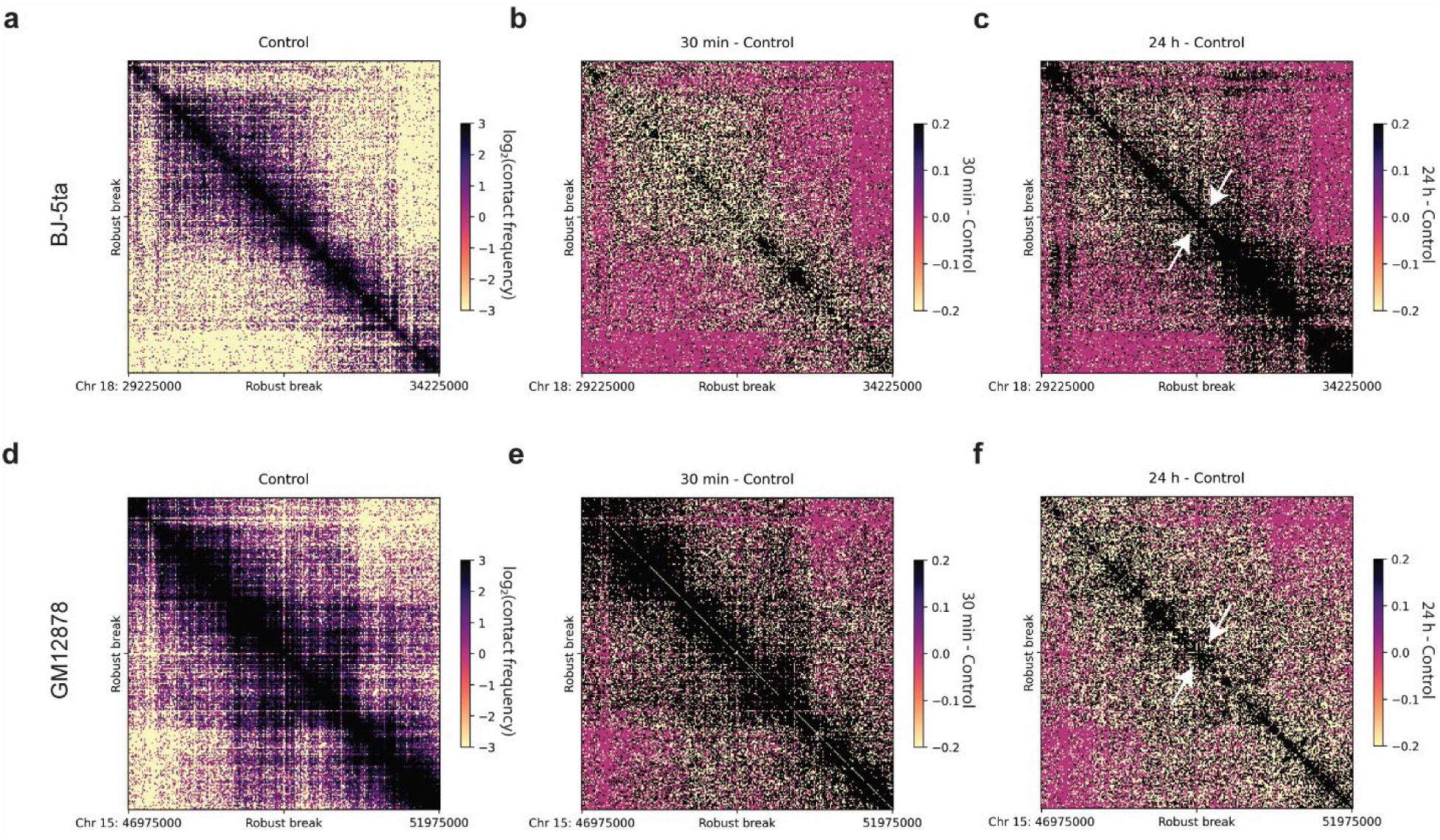
Loop extrusion at robust DSB 24 h after IR a, **d.** Hi-C contact matrix centered on a selected robust DSB in nonirradiated (control) BJ-5ta (**a**) and GM12878 (**d**) (50-kb resolution, 5-Mb window). **b, e.** Hi-C contact difference matrix (30 min after X-ray – Control) centered on the robust DSB in BJ-5ta (**b**) and GM12878 (**e**) (50-kb resolution, 5-Mb window). **c, f.** Hi-C contact difference matrix (24 h after X-ray - Control) centered on the robust DSB in BJ-5ta (**c**) and GM12878 (**f**) (50-kb resolution, 5-Mb window).

## Discussion

Recent studies have mapped the restriction enzyme-induced and UV-induced DNA breaks and depicted the altered landscape of 3D genome organization at break sites^10,12,13^. While we previously characterized the effects of irradiation on 3D genome structure, how these changes correlate with the landscape of radiation-induced DNA breaks needed further study^14^. In this study, we focused on how the initial 3D genome organization impacts the distribution of ionizing radiation-induced DSBs and how local genome structures are reshaped by DNA break sites. Accordingly, we have performed a sensitive method (END-seq) to map the high frequency DSBs after exposure to 5 Gy IR and aligned to our Hi-C data^14^.

It has long been characterized that ionizing radiation tends to induce DNA damage randomly distributed across the whole genome. In line with this expectation, our sequenced sites of damage are distributed throughout chromosomes and do not produce the kind of focused peaks observed after AsiSI induction or etoposide. However, this widespread nature of the damage does not preclude the existence of patterns. Investigations of chromosomal aberrations have shown that breakpoints preferentially occur in Giemsa-labeled regions (G-bands) in chromosomes in mammalian lymphocytes, corresponding to euchromatin^16–18,57^. Although IR-induced breaks were enriched in heterochromatin in plant^58,59^, these data also demonstrated that the distribution of radiation-induced DNA breaks are non-random and influenced by the existing chromatin structure. Moreover, studies report that a reduction of histone proteins results in an increase in DNA damage^19^. A sequencing-based method (BLISS) has shown the consistent result that regions with lower nucleosomal occupancy are more susceptible to DNA double-strand breaks^20^. In addition, with staining of DSB foci, multiple studies revealed that heterochromatin, corresponding to condensed genomic regions, protects DNA from radiation-induced DNA damage^21–23^. Similar results have been reported by a computational simulation model^24^. Thus, these data indicate that DSB probabilities might be influenced by the DNA accessibility. To illustrate how chromatin accessibility associates with the genomic distribution of DSBs after IR exposure, we conducted a multiscale analysis of DSB distribution across hierarchical levels of chromatin organization, including chromosome territories, A/B compartments, and TADs. At the chromosome scale, DNA breaks occur more frequently in open and gene-rich chromosomes (e.g., chr 19) compared to condensed and gene-poor chromosomes (e.g., chr 18). At the compartment level, transcriptionally active A compartments exhibit more frequent DNA damage than transcriptionally inactive B compartments. At the TAD level, TAD boundary regions, which are more accessible to insulator proteins and transcriptional machinery, are more prone to DNA damage than internal TAD regions.

Across our analyses, comparison of the number of robust break bins and break frequency revealed an inverse relationship across the chromosome structure hierarchy. Higher break frequency was accompanied by a lower number of robust break locations in accessible regions (chromosome 19, p arm of chromosome X, and A compartments), whereas lower break frequency was associated with a higher number of individual sites of robust break in condensed regions (chromosome 18, q arm of chromosome X, B compartments). These findings suggest that open chromatin may be more susceptible to damage in a larger fraction of cells, but at a more focused, limited set of genomic loci, while condensed chromatin may experience less frequent breaks at more independent locations, perhaps reflecting stochastic damage followed by slower repair kinetics in this type of chromatin.

Our results also showed that an unusually large number of robust breaks occur on chromosome X in both cell types. This may indicate that repair is inefficient on chromosome X. The inactive X chromosome is condensed and transcriptionally silent in female cells and chromosome X and Y lack a homologue in male cells. Given that DNA repair efficiency is impaired by heterochromatin^60–62^ and the absence of a homologous template, we propose that DNA damage is accumulated and persistent on chromosome X. In lymphoblasts, robust breaks on chromosome X were unevenly distributed, with a significantly higher recurrence on the p arm despite a greater total number of breaks on the q arm. This asymmetry corresponded with a higher gene expression density on the p arm. It has been shown that the majority of genes escaping from X-chromosome inactivation are distributed in the short arm of chromosome X and they mainly locate at the proximal or the distal third of the short arm^63–66^. This suggests that transcriptional activity may contribute to site-specific susceptibility. While only a subset of DNA breaks align with loci of known escaping genes, this hypothesis requires further investigation to assess whether the remaining break-prone regions may escape X-chromosome inactivation in this particular cell type, leading them to be more sensitive to radiation.

In line with the difference of DSBs distribution on chromosome X in fibroblasts and lymphoblasts, we found that the same dose of IR induces distinct number of robust breaks in each chromosome across different cell types. This observation suggests that differences in nuclear architecture may contribute to cell-type specific DNA damage patterns. Our simulations on the arrangements of chromosomes using Hi-C data by G-NOME represented the divergent cell nucleus model^32^. Compared to the flattened nucleus model in fibroblasts, lymphoblast nucleus shows a spherical configuration, with most chromosomes positioned near the nuclear periphery, causing a more uniform distribution of robust breaks across chromosomes. These findings highlight the importance of initial 3D genome organization in shaping the susceptibility of individual chromosomes to DNA damage. Meanwhile, we observed cell-type specific alterations in 3D genome organization after IR. Compared to fibroblasts, lymphoblasts showed fewer compartment shifts associated with robust breaks and weaker stripe-like patterns flanking DSB sites. While previous work has observed compartment shifts after DNA damage in fibroblasts, compartment shifts after IR have not been previously measured in lymphoblasts. We hypothesize that these differences may be caused by the cell cycle state, as fibroblasts were mostly in G1 phase, while lymphoblasts were unsynchronized.

While phosphorylation of H2AX is widely used to monitor the DNA damage response, γH2AX can spread over neighboring chromatin for 1-2 Mb and appear as diffuse nuclear foci, indicating a lack of resolution of precisely identify DSB sites. With the advent of genome-wide DSBs mapping technology END-seq, it is possible to detect AsiSI-induced DSBs at annotated sites at nucleotide resolution^37^. However, IR-induced DNA damage does not occur at reproducible single nucleotide positions in all cells. IR induces DNA damage through directly interacting with DNA molecules and indirectly ionizing water to generate free radicals and ROS which can damage DNA, lipid, and proteins, leading to the complexity of IR-induced DSBs^15^. Compared to DSBs which are consistently and faithfully cleaved by AsiSI in each cell, it is much more rare for IR-induced DSBs to occur at the same location across cells in the population. Therefore, although we identified thousands of regions enriched for IR-induced frequent DSBs, the current resolution is limited to 50 kb. Further studies are needed to determine whether there exist any more precise hotspots of DNA damage after irradiation.

Compared to the BLESS and BLISS methods of mapping DSBs, END-seq embeds native cells in agarose gel plugs, thereby protecting genomic DNA from artifacts introduced by formaldehyde fixation and mechanical processing. However, endogenous DNA repair machinery may become activated prior to lysis, potentially affecting the detection of certain types of DNA breaks. DSBs are primarily repaired through two major pathways: non-homologous end joining (NHEJ) and homologous recombination (HR), which differ in their repair fidelity and regulation across cell cycle^67^. It has been shown that proteins involved in DSB signaling and scaffolding, such as 53BP1, Ku70/80 and DNA-PKcs, are recruited within seconds to 30 minutes to DSB sites^68–71^. Moreover, real-time estimation of DNA repair protein dynamics has revealed the recruitment of XRCC4 to DSBs within 30 minutes, although it is slower than Ku70/80^71^. This suggests a fast response and repair by the NHEJ pathway, decreasing the ability of END-seq to detect these lesions. Concurrently, we observed an increasing enrichment of frequent breaks switching from A to B compartments in fibroblasts, similar to the movement seen in transcription-coupled DSBs which are repaired by HR at the nuclear envelope^9^. An optimized experiment is required to investigate DSBs repaired by NHEJ in the future.

Overall, our study sheds light on the distribution of robust radiation-induced DNA breaks, which have higher frequency in the open and transcriptionally active regions, as well as the alterations of 3D genome organization at these sites in both fibroblasts and lymphoblasts. Our results demonstrate that END-seq can detect the differential patterns of radiation-induced DNA damage across various cell types, providing a means to assess how pre-treatment conditions influence the distribution of genomic susceptibility.

## Methods

### Cell culture and treatments

BJ-5ta (RRID: CVCL_6573) and GM12878 (RRID: CVCL_7526) cells were purchased from ATCC (CRL-4001) and Coriell respectively. The U2OS-ER-AsiSI cell line was kindly provided by Gaëlle Legube (Centre de Biologie Intégrative). The BJ-5ta-ER-AsiSI cell line was kindly provided by Fabrizio d’Adda di Fagagna (Istituto Fondazione di Oncologia Molecolare ETS). BJ-5ta cells were cultured in a complete medium (4:1 mixture of Dulbecco’s Modified Eagle’s Medium (DMEM, Corning, 10-013-CV) and Medium 199 (M199, Gibco, 11150-059), supplemented with 10% FBS, 1% Pen-Strep, 1% L-glutamine and 0.01 mg/ml hygromycin B) at 37°C supplied with 5% CO_2_. GM12878 cells were cultured in complete Roswell Park Memorial Institute Medium 1640 (Gibco, 11875-093; 15% FBS, 1% Pen-Strep, 1% L-glutamine and 0.01 mg/ml hygromycin B) at 37°C supplied with 5% CO_2_. BJ-5ta and GM12878 cells were cultured and passaged as previously described. For irradiation experiments, BJ-5ta adhesion cells were grown to confluency, and GM12878 suspension cells were grown to a density of 1 × 10^6^ cells per 1 mL medium prior to X-ray exposure. U2OS-ER-AsiSI cells were cultured in a complete DMEM GlutaMAX medium (Gibco, 10566-024), supplemented with 10% FBS, 1% Pen-Strep, 1 mM sodium pyruvate and 1 µg/mL puromycin) at 37°C supplied with 5% CO_2_. BJ-5ta-ER-AsiSI cells were cultured in a complete medium (4:1 mixture of DMEM and M199, supplemented with 10% FBS, 1% Pen-Strep, 1% L-glutamine, 0.01 mg/ml hygromycin B and 1 µg/mL puromycin) at 37°C supplied with 5% CO_2_. U2OS-ER-AsiSI and BJ-5ta-ER-AsiSI cells were cultured and passaged with puromycin selection. For DNA damage induction, both cells were seeded without puromycin treatment and treated with 300 nM 4-hydroxytamoxifen (4OHT) for 4 h and 6 h, respectively. 4OHT (Sigma, H7904) was dissolved in dimethyl sulfoxide (4-X, ATCC) and stored at −80°C with a stock concentration of 10 mM. All cells were verified to be negative for mycoplasma.

### *X-* ray irradiation

Cells were irradiated using a RS 2000 X-ray Irradiator (RadSource). A 2 Gy/min dose rate was used to deliver a total of 5 Gy X-rays (160 kV; 25 mA). After irradiation, cells were immediately processed for the subsequent experiments.

### END-seq

END-seq was performed in U2OS-ER-AsiSI, BJ-5ta-ER-AsiSI, BJ-5ta and GM12878 cells as described^29^ for two biological replicates. Cells were immediately collected and embedded into an agarose gel plug (Bio-Rad Laboratories) after exposure to 5 Gy X-rays (BJ-5ta and GM12878 cells) or 4OHT treatment (U2OS-ER-AsiSI and BJ-5ta-ER-AsiSI cells). Exonuclease VII and exonuclease T (New England Biolabs) were used to process and cleave damaged ends. The biotinylated Illumina adapter 1 was added and ligated to the 3’ dA-tailing blunted ends. After removing extra adapter 1, the agarose plug was melted and released DNA was extracted and sheared to a target medium size of 175 bp a Covaris sonicator (Covaris, M220). The cycle setting on sonicator was peak power 50, duty 14, cycles/burst 200, 420 s, 4-7 °C.

Biotin-labeled DNA was pulled down using streptavidin coated beads (Dynabeads MyOne C1) and the new ends created by sonication were blunted and dA-tailed, allowing ligation of Illumina adaptor 2. The hairpins of adaptors were digested with USER enzymes (NEB). DNA was amplified with primers including unique barcodes. Pooled PCR libraries were sequenced at the University of Colorado Anschutz Medical Campus Genomics Core on an Illumina NovaSeq 6000 platform.

### END-seq data processing

END-seq data were aligned on human reference genome GRCh37 (hg19) using Bowtie2 and processed into counts per bin using CUT&RUNTools2.0^72,73^. Raw data for each aligned dataset were normalized in each genomic bin by dividing the counts by total read count for each sample (counts per million) and exported as bigwig (file format) for further analysis.

### Analysis of DSBs-induced via AsiSI

To identify which of the 1211 AsiSI sites were actively cleaved in U2OS-ER-AsiSI and BJ-5ta-ER-AsiSI cells, bigwig files created as described above were uploaded to the Galaxy web platform and processed with computeMatrix tool to quantify the average END-seq signal within 1 kb windows centered on each site^74,75^. The average END-seq signal was ploted in profile and heatmap by using plotProfile and plotHeatmap tools^75^. AsiSI sites in U2OS-ER-AsiSI cells were defined as robust DSB sites if the average END-seq signal within 0.6 kb windows centered on each site was greater than 0.4 in both replicates.

### Analysis of DSBs induced via X-ray

To determine the distribution of radiation-induced DSBs in BJ-5ta and GM12878 cells after exposure to 5 Gy X-rays, bigwig files including both X-ray treated and untreated control conditions, created as described above were used in the following pipelines. Read counts were aggregated into 10 kb genomic bins. Gap annotations, including centromeres and telomeres and known artifacts, were downloaded from the UCSC Genome Browser and extended by 1 Mb on each side to account for edge effects and low-complexity sequences.

A threshold to detect artifactually high signal outside of known gaps was defined as the 99.9th percentile of signal intensities in non-gap regions. Bins within the extended gap regions exceeding this threshold were flagged in each replicate and treatment condition (R1-X-ray, R2-X-ray, R1-control, R2-control). To ensure consistency and avoid replicate-specific artifacts, the flagged bin sets were merged to generate a unified exclusion list for each cell type. This list was applied to all replicates to remove bins with potential mapping artifacts.

Filtered 10 kb bins were aggregated into 50 kb bins to improve signal robustness. For each 50 kb bin, the log₂ ratio of signal intensity between X-ray treated and control conditions was calculated. Bins with log₂ ratios greater than 1 in both replicates were defined as robust double-strand break (DSB) regions, reflecting at least a twofold increase in signal relative to control. To avoid division by zero or inflation from low counts, a pseudocount of 0.00001 was added to all signal intensity values prior to ratio calculation. This threshold 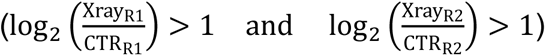 was selected to capture regions with consistent enrichment of DSBs occurring after X-ray across replicates.

Throughout the remaining analyses, DSBs were quantified in two ways: Break Counts = total number of 50 kb bins classified as “robust breaks” by criteria above (log₂ ratio > 1 in both replicates). Break Frequency = sum of signal intensity for the X-ray treated condition within bins classified as “robust breaks”, computed separately for each replicate (R1 and R2).

### Correlation Between Chromosome Length and DSB Counts

Chromosome length information (hg19 assembly) was obtained from the UCSC Genome Browser. The relationship between chromosome length and the number of robust DSB regions was evaluated by first normalizing total chromosome length, subtracting the length of gap-proximal regions excluded during signal filtering.

For BJ-5ta cells, DSB counts on autosomes were divided by two, while counts in sex chromosomes (chr X and chr Y) were left unchanged. For GM12878 cells, DSB counts on all chromosomes were divided by two. This normalization reflects the effective haploid genome content, enabling comparison at the single-copy level.

Spearman correlation coefficients were calculated between normalized chromosome lengths and Break Counts per chromosome.

### Hi-C data analysis

Raw Hi-C data reads were available and accessible in Gene Expression Omnibus (GEO) under accession number GSE136899. The data analysis was performed same as the previous study^14^.

Briefly, compartment and TAD boundary analysis were performed using matrix2compartment.pl and matrix2insulation.pl scripts in the cworld-dekker pipeline available on GitHub (https://github.com/dekkerlab/cworld-dekker), respectively. The principal component analysis was performed on 250 kb binned matrices and determined the A and B compartments. Compartment switches were identified as regions exhibiting an absolute change in PC1 values greater than 0.01 compared to the control sample.

TAD boundaries were called on 40 kb binned matrices and insulation score was calculated by InsulationScore method^76^.

### Integration of Hi-C and END-seq data

#### Assignment of A/B Compartments and Compartment Shift Analysis

Robust DSB bins (50 kb size) were assigned compartment identity based on Hi-C data described above using PyRanges^77^. Enrichment analysis of robust DSBs located in A or B compartments were calculated separately for each cell type.

Compartment scores from Principal Components Analysis were compared between control and X-ray treated conditions (30 min and 24 h). Regions showing A-to-B or B-to-A transitions with an eigenvector value change > 0.01 were defined as compartment switches and intersected with robust DSBs. The enrichment of robust DSBs in switching compartments was assessed using Fisher’s exact tests. For both control vs. 30 min and control vs. 24 h comparisons, contingency tables were constructed to compare bins with or without DSBs across regions with or without compartment flips.

#### TAD Boundary Analysis

TAD boundary coordinates for BJ-5ta and GM12878 were intersected with robust DSBs using PyRanges^77^. For each chromosome, total break counts were computed and partitioned into boundary and non-boundary groups for each chromosome. Effective lengths of boundary regions (summed boundary intervals) and non-boundary regions (chromosome length minus boundary length and excluded gap regions) were calculated and used to normalize DSB counts to the megabase level.

Box plots were generated to compare break counts between boundary and non-boundary regions. Statistical significance was assessed using the Mann–Whitney U test.

#### Position analysis

The radial position of each chromosome in the nucleus was obtained from previously published data^78^. Robust DSB counts were summed per chromosome, with counts on autosomes divided by two as described above.

Scatter plots were generated to visualize the relationship between chromosome position and normalized DSB frequency. Spearman correlation was computed to assess the association.

#### Chromosome X Arm-Level Analysis

RNA-seq data for BJ-5ta and GM12878 were obtained from the Gene Expression Omnibus (GEO) under accession numbers GSE247831^45^ and GSM5664127^46^. Cytoband annotations were downloaded from the UCSC Genome Browser to define chromosomal arms, using the centromere coordinates to identify the p and q arms of chromosome X. Robust DSBs were then assigned to either arm based on genomic coordinates.

Box plots were generated to visualize the distribution of DSB frequency across the two arms. Statistical significance was assessed using the Mann–Whitney U test. To evaluate transcriptomic differences, gene counts on each arm were summed and normalized by effective arm length to obtain counts per megabase. Bar plots were generated to display the normalized total expression on the p and q arms.

#### Pileup Analysis of Hi-C Signal at DSB Sites

To examine local chromatin architecture around robust DSBs, pileup Hi-C maps were generated using cooltools^79^. DSBs were first ranked by the sum of X-ray–minus–control End-seq signals across replicates: (Xray_R1 − CTR_R1) + (Xray_R2 − CTR_R2). The midpoint of each DSB region was used as the anchor, with a flanking window of ±25 kb applied to extract the local contact matrix.

For each cell type, pileup maps were generated separately for control and X-ray treated samples. Differential maps were computed by subtracting the control signal from the treated condition to highlight structural changes associated with DSB induction.

## Data Availability

All new sequence data and processed data files for END-seq data are available at GEO Accession number GSE306338.

## Code Availability

Analysis code is available upon request.

## Funding

This work was supported by the National Institutes of Health NIGMS grant R35GM133557, Department of Energy award DE-SC0025274, and UT-ORNL Science Alliance StART award to R.P.M. H.L. was supported by a University of Tennessee-Oak Ridge Innovation Institute GATE fellowship.

## Acknowledgements

The U2OS-ER-AsiSI cell line was kindly provided by Gaëlle Legube (Centre de Biologie Intégrative). The BJ-5ta-ER-AsiSI cell line was kindly provided by Fabrizio d’Adda di Fagagna (Istituto Fondazione di Oncologia Molecolare ETS). We thank Dr. Andres Canela, Brandon Estrem, and Jianbin Wang for advice and guidance on END-seq experiments.

## Author Contributions

The study was designed by H.L and R.P.M. H.L. performed all experiments. T.X. performed computational data integration and analysis. Figures and graphs were made by H.L. and T.X. H.L., T.X., and R.P.M. wrote the manuscript.

## Competing Interests

The authors declare no competing interests.

## Notes

https://www.ncbi.nlm.nih.gov/geo/query/acc.cgi?acc=GSE306338

### Competing Interest Statement

The authors have declared no competing interest.

