## Supplementary Figures for "The multiscale distribution of radiation-induced DNA damage and its impact on local genome structure"

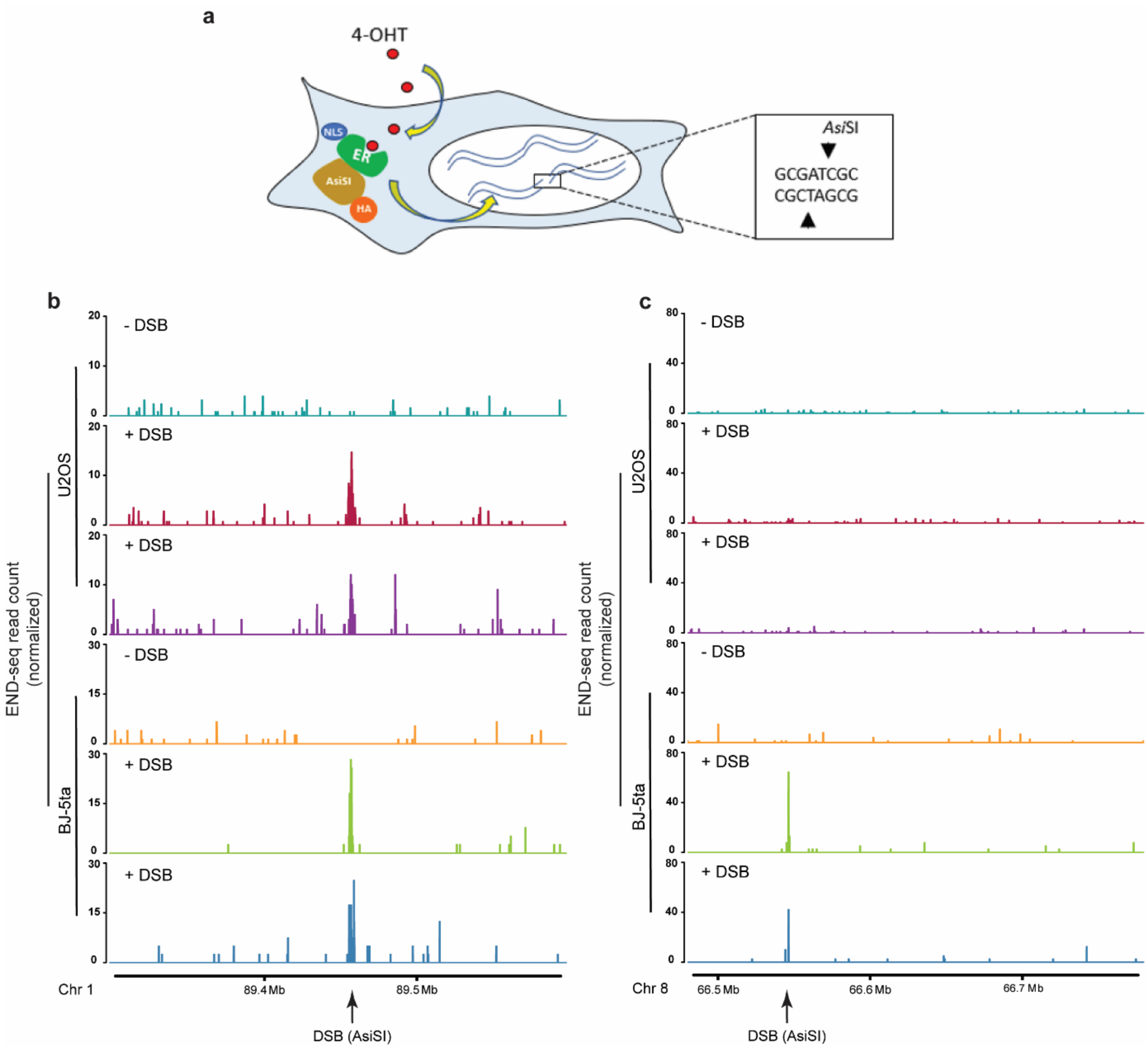


**Supplementary Figure 1: Genome-wide mapping of DSBs by END-seq in DIvA cell lines a.** Schematic of the DIvA cell line. Restriction enzyme AsiSI is fused to a modified oestrogen receptor, which responds to 4OHT. AsiSI-ER fusion protein is imported to the nucleus upon 4OHT treatment. AsiSI digests an 8 bp target sequence (GCGATCGC) across the genome and generates DSBs. **b.** Stably expressed AsiSI-ER cell lines, U2OS (top) and BJ-5ta (bottom), were treated with 4-OHT to induce DSBs at specific genomic loci. END-seq signal shows reproducible peaks in two replicates after AsiSI induction at a known AsiSI cut site (arrow) **c.** An example of a cell type specific DNA damage site where END-seq signal shows consistent DSBs at a known AsiSI site (arrow) in BJ-5ta-ER-AsiSI cells but not U2OS-ER-AsiSI cells.

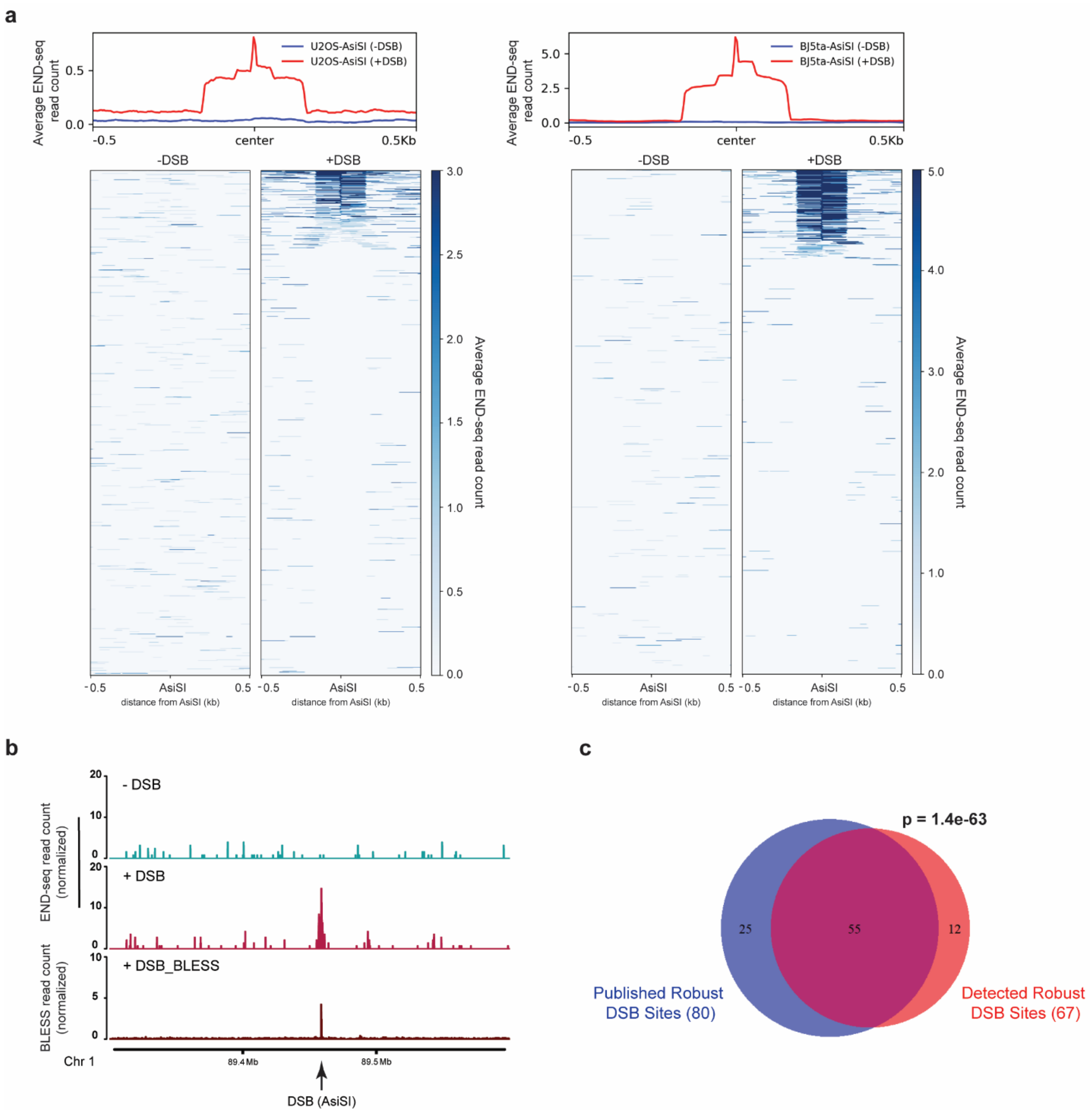


**Supplementary Figure 2: Identification of robust AsiSI-induced DSBs a.** Average END-seq signal profile and heatmap for the 1211 predicted AsiSI sites in the human genome in U2OS-ER-AsiSI (left) and BJ-5ta-ER-AsiSI (right) cells before (-DSB, blue line) and after (+DSB, red line) induction. **b.** END-seq data compared with previously published BLESS (+DSB) tracks (Clouaire et al., 2018) on a 300 kb window around an AsiSI site (arrow) in U2OS-ER-AsiSI cells. **c.** Venn diagram depicting the robust DSBs at predicted AsiSI sites detected by BLESS (80) and END-seq (67) in U2OS-ER-AsiSI cells. A Fisher's exact test determined that the overlap between DSBs detected in both BLESS and END-seq method is statistically significant.

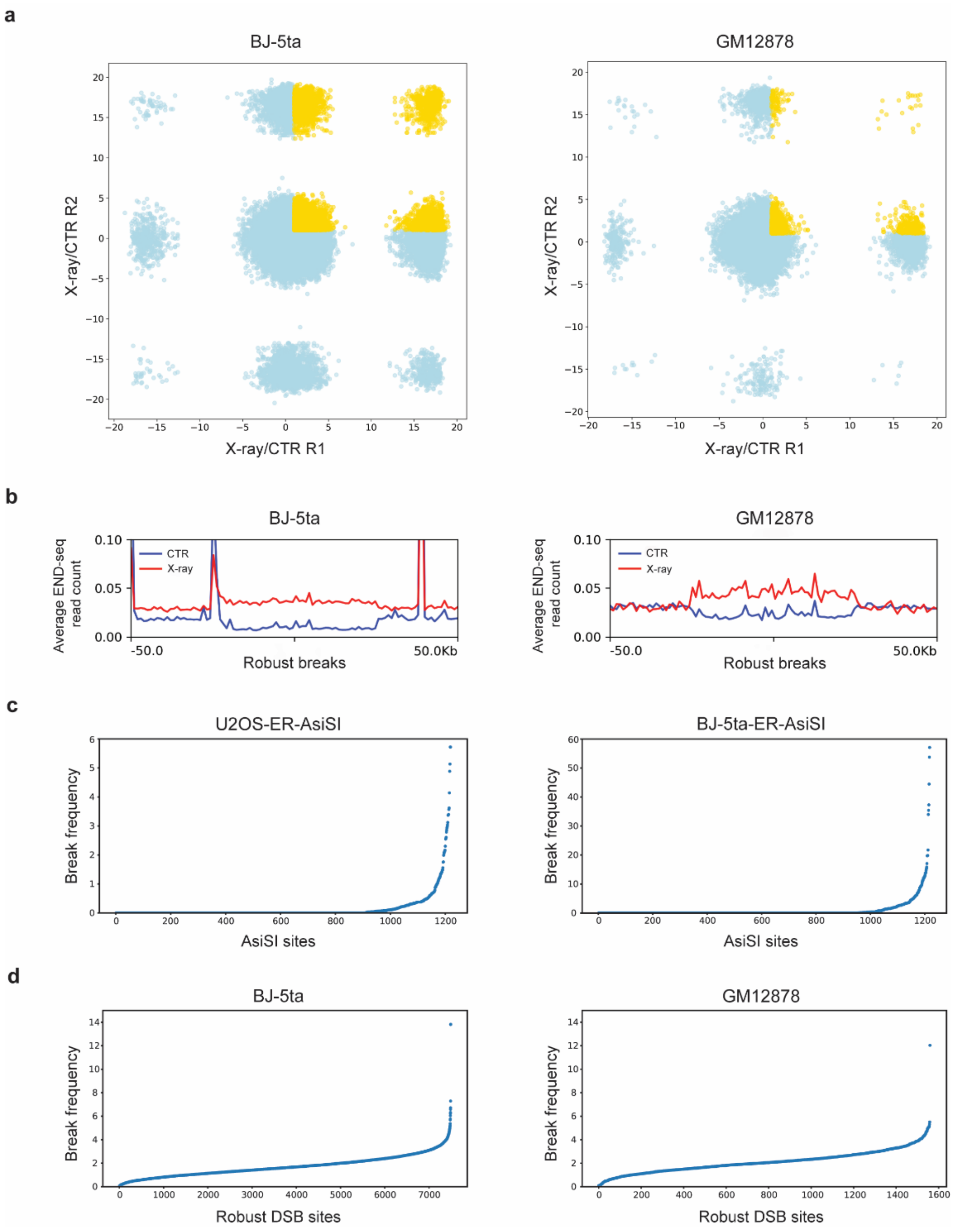


**Supplementary Figure 3: Identification of robust breaks after IR a.** Scatter plots of two replicates showing the log2 ratio of END-seq signal between X-ray and Control in BJ-5ta (left) and GM12878 (right) cells. Shared breaks whose log ratios are greater than 1 in both replicates are labeled in yellow. **b.** Average break frequency profile (from biological replicate 1) centered on the 50 kb robust break site locations in the human genome in BJ-5ta (left) and GM12878 (right) cells without (CTR, blue line) and with (X-ray, red line) X-ray exposure. Break frequency is binned at 1 kb. **c.** Dotplot representing END-seq signal within a 1 kb window centered on 1211 predicted AsiSI sites in the human genome in U2OS-ER-AsiSI (left) and BJ-5ta-ER-AsiSI (right) cells. Sites are ranked in ascending order by increasing break frequency signal. **d.** Dotplot representing END-seq signal within 50 kb windows for the robust break sites in the human genome in BJ-5ta (left) and GM12878 (right) cells. Sites are ranked in ascending order by increasing signal.

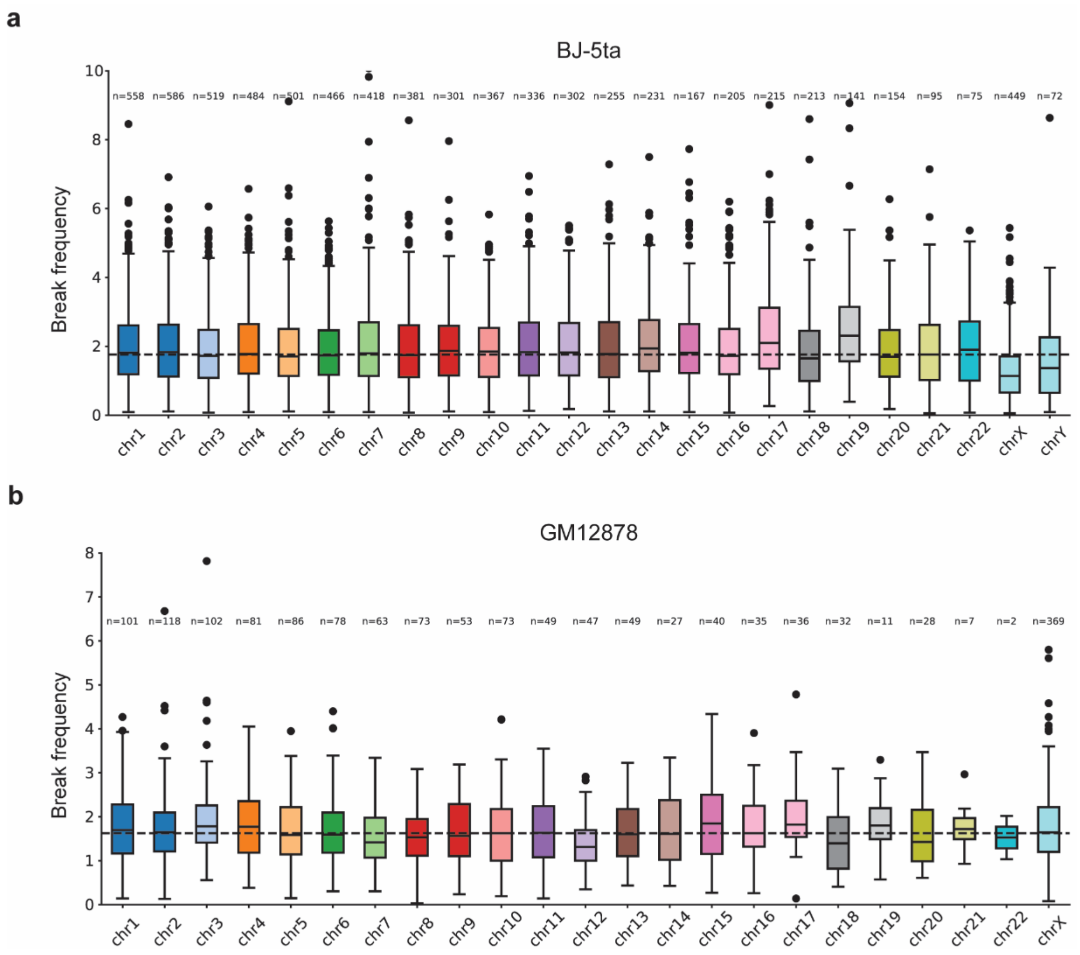


**Supplementary Figure 4: Robust break frequency measurements across chromosomes in a separate biological replicate** Boxplots representing the break frequency in 50 kb windows along chromosomes in BJ-5ta (**a**) and GM1287 (**b**) cells. Boxes are colored by chromosomes.

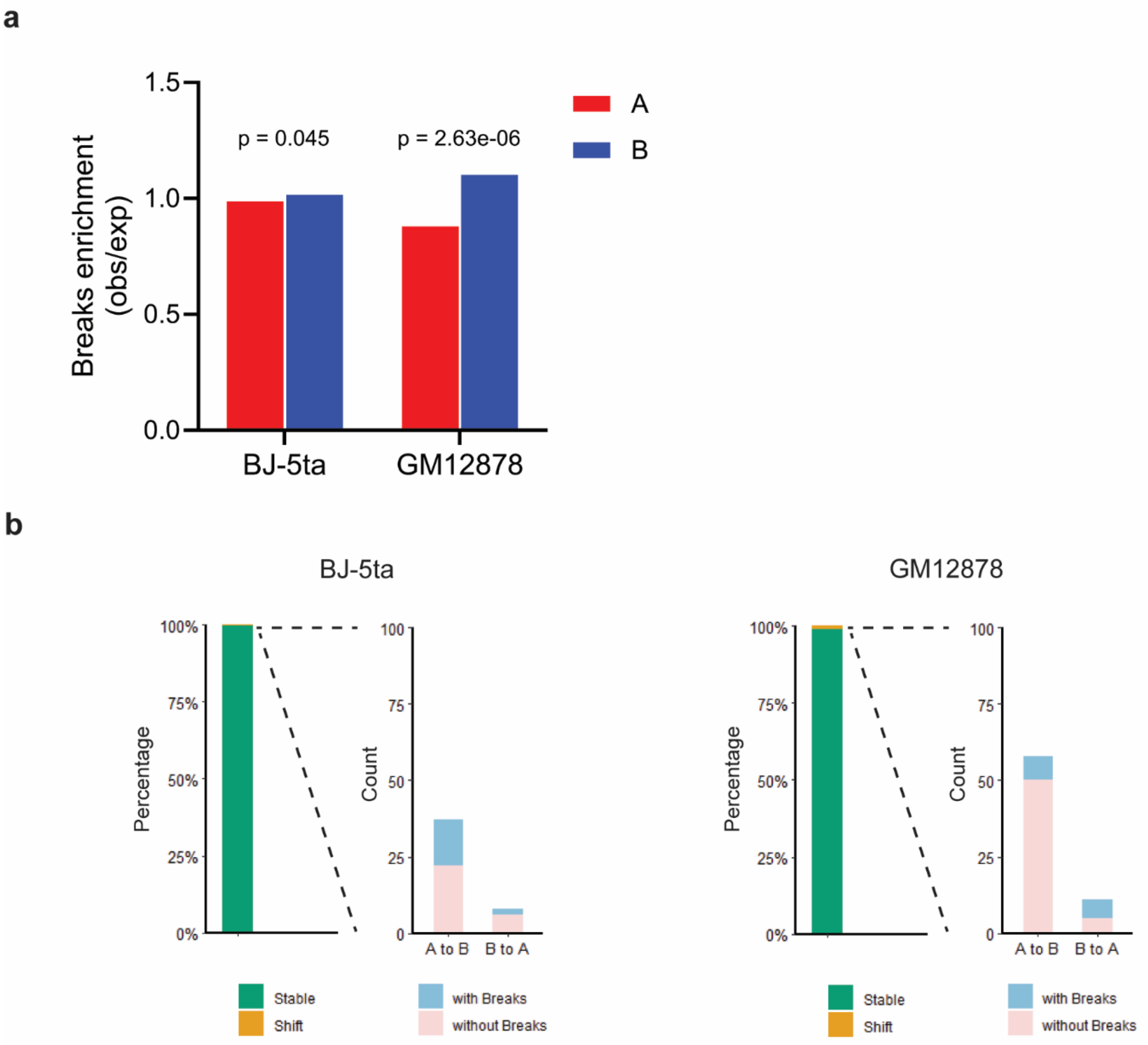


**Supplementary Figure 5: Robust break enrichment analysis in A/B compartments and proportion of break-associated shifts at 24h after IR a.** Bar plots showing the enrichment analysis of robust breaks (obs/exp) located within A (red) and B (blue) compartments in BJ-5ta (left) and GM12878 (right) cells. **b.** Bar plots showing the counts of 250 kb bins with (sky blue) and without (rose) robust breaks within compartments that shifted between control and 24 h post-irradiation, including A to B and B to A, in BJ-5ta (left) and GM12878 (right) cells.
